# *Arabidopsis* stomatal polarity protein BASL mediates distinct processes before and after cell division to coordinate cell size and fate asymmetries

**DOI:** 10.1101/2021.06.18.448880

**Authors:** Yan Gong, Julien Alassimone, Andrew Muroyama, Gabriel Amador, Rachel Varnau, Ao Liu, Dominique C. Bergmann

## Abstract

In many land plants, asymmetric cell divisions (ACDs) create and pattern differentiated cell types on the leaf surface. In the Arabidopsis stomatal lineage, *BREAKING OF ASYMMETRY IN THE STOMATAL LINEAGE* (*BASL*) regulates multiple aspects of ACD including division plane placement and cell fate enforcement. Polarized subcellular localization of BASL is initiated before the ACD and persists for many hours after the division in one of the two daughters. Untangling the respective contributions of polarized BASL before and after division is essential to gain a better understanding of its roles in regulating stomatal lineage ACDs and to uncover the rules that guide leaf pattern. Here we combine quantitative imaging and lineage tracking with genetic tools that provide temporally-restricted BASL expression. We find that pre-division BASL is required for division orientation, whereas BASL polarity post-division ensures proper cell fate commitment. These genetic manipulations allowed us to uncouple daughter-cell size asymmetry from polarity crescent inheritance, revealing independent effects of these two asymmetries on subsequent cell behavior. Finally, we show that there is coordination between the division frequencies of sister cells produced by ACDs, and this coupling requires BASL as an effector of peptide signaling.

## INTRODUCTION

Cell polarity underlies many of the asymmetric and oriented cell divisions that create daughter cells with distinct fates (Drubin & Nelson, 1996; Muroyama & Bergmann, 2019). In both plants and animals, cell polarity frequently manifests as protein enrichment in specific subdomains of the cell, often associated with the plasma membrane. During asymmetric cell division (ACD) in metazoans, spatially restricted “polarity proteins” and their interaction partners orient cell division planes and recruit cell fate determinants to ensure asymmetric inheritance in daughter cells (Morin & Bellaiche, 2011). The general picture emerging from several decades of genetic analysis and protein-protein interaction studies is that polarity proteins act as scaffolds before division, both by recruiting clients that can guide spindle orientation and by creating environments that exclude proteins (e.g. cell fate determinants) from certain regions of the mother cell (Morin & Bellaiche, 2011).

Despite similar needs for cell polarity to orchestrate cellular and developmental behaviors, plants appear to employ a different suite of polarity proteins from those used in animals. One such protein, BREAKING OF ASYMMETRY IN THE STOMATAL LINEAGE (BASL), plays a central role in directing nuclear migrations to orient division planes and scaffolding intracellular signaling cascades to solidify differential cell fates in the *Arabidopsis* stomatal lineage (Dong et al., 2009; Muroyama et al., 2020; Zhang et al., 2016; Zhang et al., 2015). The stomatal lineage features asymmetric, oriented, and self-renewing divisions (Figure 1A) and has emerged as a powerful model for investigations of how cellular polarity specifies developmental patterns in plants.

**Figure 1.**
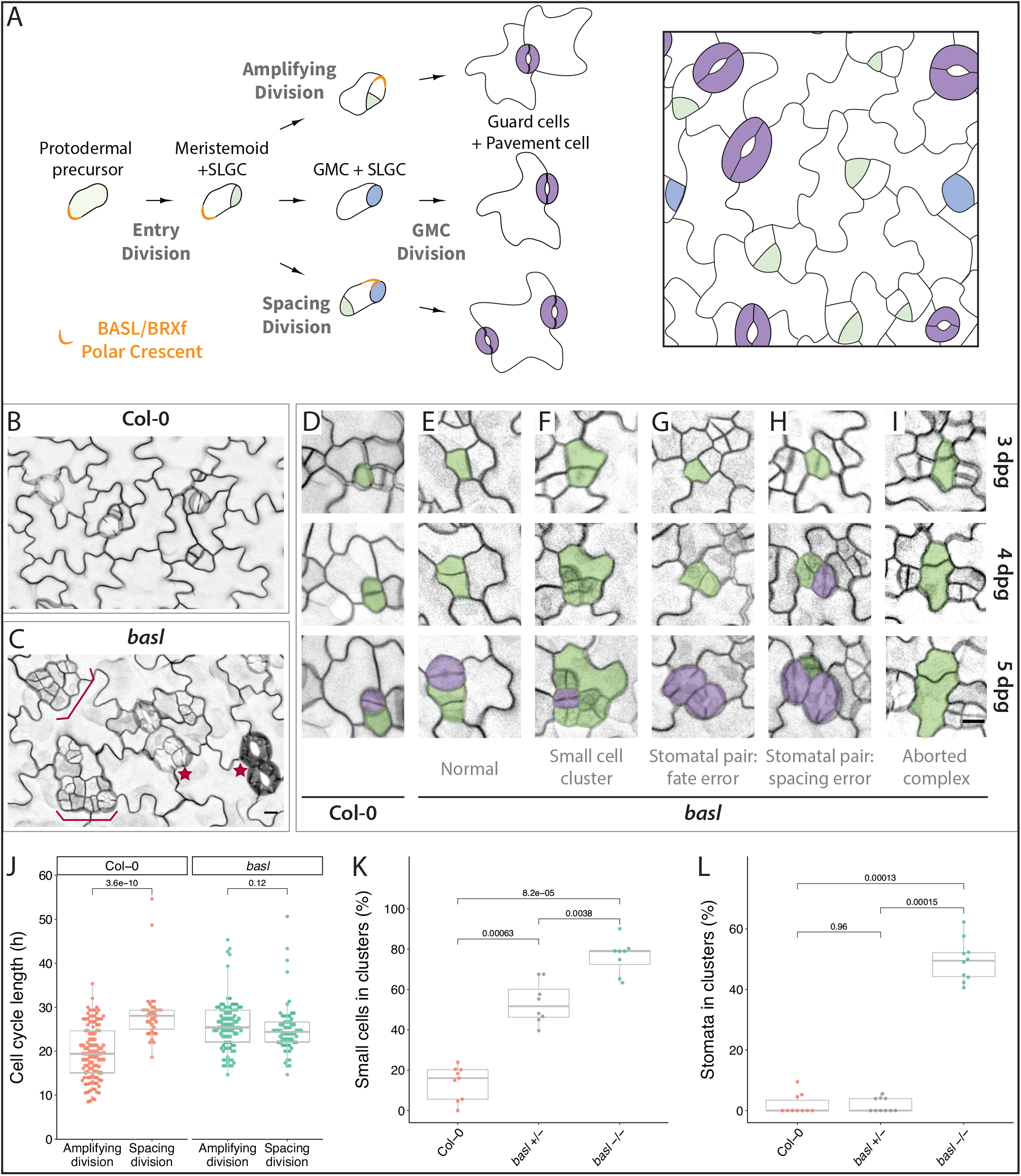
Lineage tracing and quantitative analysis reveals multiple roles for BASL in the *Arabidopsis* stomatal lineage. (A) Cartoon of stomatal lineage cell divisions with associated fate outcomes (left) and their arrangements on the leaf epidermis (right). Protodermal precursor cells enter the lineage and undergo asymmetric “entry” divisions producing a smaller meristemoid (green) and a larger stomatal lineage ground cell (SLGC, white) daughter. Before division, BASL and partner proteins form a polar crescent at the cell cortex (orange); this is inherited by the SLGC through the division. Both meristemoids and SLGCs can undergo additional ACD(s), termed amplifying divisions and spacing divisions, respectively. Alternatively, the SLGC can undergo terminal differentiation into a pavement cell and the meristemoid can transition into a guard mother cell (GMC, blue), undergo one symmetric division, and produce paired guard cells (purple). At any given time during leaf development, dispersed stomatal lineage cells are actively initiating, dividing, and differentiating. (B-C) Confocal images of the abaxial epidermis of 5 dpg cotyledons from Col-0 (B) and *basl* (C) seedlings. Cell outlines are visualized by plasma membrane reporters *pATML1::RCI2A-mCherry* (Col-0) or *p35S::PIP2A-RFP* (*basl*). Braces and stars indicate clustered small cells and stomata, respectively. (D-I) Examples of lineage-traced stomatal lineage cells from Col-0 (D) and *basl* (E-I) abaxial cotyledons from 3 dpg to 5 dpg, showing the origins of normally patterned cell types (D-E), small cell clusters (F), stomatal pairs (G-H), and aborted complexes (I). Early stomatal lineage cells and stomata are false colored in green and purple, respectively. (J) Cell cycle lengths among stomatal lineage cells from Col-0 (n=147) and *basl* (n=151) seedlings. Spacing and amplifying refer to divisions of the large and smaller daughter cell, respectively, that were created by the previous division. (K) Quantification of small cell clusters in the 5 dpg abaxial cotyledon epidermis (n=8 388*388 μm2 fields/genotype) The Bonferroni correction was applied to reduce Type I error, and the corrected p-value threshold for rejecting the null hypothesis (no difference between the mean of the compared groups) was calculated to be 0.017 (0.05/3). Any p-values below this corrected threshold were deemed significant. (L) Quantification of clustered stomata in 14 dpg abaxial cotyledon epidermis of the same genetic background in (K) (n=10 623*467 μm2 fields/genotype, Bonferroni corrected p-value threshold = 0.017). All p-values are calculated by Mann-Whitney test. Scale bars, 10 μm.

A signature of stomatal lineage development, and a feature it shares with other plant developmental systems, is flexibility and tunability. The developmental trajectories of stomatal precursors are not fixed, and can be tuned by nutritional, hormonal, and environmental cues (Engineer et al., 2014; Gong, Alassimone, et al., 2021; Han et al., 2020; Lau et al., 2018; Lee & Bergmann, 2019; Vaten et al., 2018; Wang et al., 2021). BASL polarity has been linked to this flexibility in several ways: BASL is employed to enforce cell fate commitment after ACDs (Dong et al., 2009; Muroyama et al., 2020; Zhang et al., 2016; Zhang et al., 2015), and recent work has also correlated the persistence of BASL after ACD with self-renewing division capacity (Gong et al., 2021). Roles for polarity post-division, particularly in regulating lineage flexibility, have been relatively understudied, in part because they are not features of the rapid and invariant cell divisions typical of invertebrate models used to investigate ACDs. In addition, there are structural differences between animal cells and walled plant cells, and these two branches of life use different cytoskeletal structures to dictate division planes and complete cytokinesis. Taken together, there remain numerous questions about the cellular and developmental mechanisms of ACDs, for which a careful examination of BASL activities in the stomatal lineage might provide new or unexpected insights.

Here, we used time-lapse imaging, lineage tracing and quantitative polarity measurements to characterize how specific asymmetries—in cell size, division rates, inherited factors, and orientation relative to landmarks—affect cell fate and the overall pattern of the leaf epidermis. By creating genetic reagents to supply BASL at discrete times over the course of asymmetric division, we show that BASL functions pre- and post-ACD are separable. Pre-division BASL is critical for division orientation while BASL polarization post-division is necessary and sufficient for daughter cell fate asymmetry Additionally, we discovered that supplying BASL post-division can uncouple cell size and polarity inheritance, revealing a previously unappreciated role for BASL as an effector of sister-cell division coordination. Together, these results reveal how separable polarity modules can be coordinated by BASL to pattern ACDs in the leaf epidermis.

## RESULTS

### Loss of BASL abolishes fate asymmetry in ACD daughter cells and disrupts patterning of the leaf epidermis

To comprehensively determine the myriad functions of BASL during epidermal patterning, we re-examined the phenotypes of *BASL* null mutants (*basl-2*, hereafter referred to as *basl*) using time-lapse imaging and whole-leaf cell lineage tracing (Gong, Alassimone, et al., 2021). With this method, we tracked the developmental progression of all epidermal cells over 48 hours (3 to 5 days post germination, dpg) in the abaxial surface of Col-0 wild-type and *basl* cotyledons expressing fluorescent reporters that mark plasma membranes. We observed that the *basl* abaxial epidermis exhibited an overall increase in cell number and numerous clusters of small cells and of mature stomata (Figure 1B-C), in agreement with previous studies (Dong et al., 2009). Our lineage tracing revealed that small cell clusters typically form from excess divisions in both the larger and smaller cells born from an ACD (Figure 1D-F). Clusters of mature stomata in the *basl* mutant arose from two independent origins. First, as reported in the original characterization of *basl* (Dong et al., 2009), both daughter cells of a cell division could become GMCs and undergo symmetric cell divisions (GMC division) to form two adjacent stomata (Figure 1G). Second, we found that stomatal pairs arose from misoriented spacing divisions, where a stomatal precursor cell is formed next to an existing stoma or GMC (Figure 1H). Lineage tracing over multiple days also revealed two previously unreported phenotypes in *basl*: (1) formation of “aborted complexes” where both daughter cells of a seemingly asymmetric cell division acquired pavement cell fate (Figure 1I), and an equalization of cell cycle timing among daughter cells resulting from ACDs (Figure 1J). Calculation of cell cycle length was especially interesting because, in contrast to many well studied systems (e.g. the *C. elegans* embryo (Arata et al., 2014; Jankele et al., 2021) or the *Arabidopsis* shoot apical meristem (Jones et al., 2017)), the smaller daughter resulting from a stomatal lineage ACD has a shorter cell cycle time than the larger daughter.

This more precise and powerful phenotypic scoring system revealed an effect of *BASL* dosage on the small cell cluster phenotype (Figure 1K), but not on the stomatal pair phenotype (Figure 1L). This has interesting implications for the relationship between these two phenotypes, as it suggests that the excessive and incorrectly patterned small cells are not always destined to become incorrectly patterned stomata. Conversely, we found that stomatal clusters can arise from activities unlinked to the production of small cell clusters (e.g., Figure 1H). Because complete *BASL* loss of function profoundly alters cell identity and division patterns, and because of the non-linear relationship between early and terminal phenotypes, we reasoned that new genetic manipulation and analysis tools would be required to precisely interrogate BASL function during the cell cycle and throughout development.

### Genetic reagents to supply BASL function pre- and post-division in asymmetrically dividing stomatal lineage cells

To precisely dissect BASL function, we designed three fluorescent BASL reporters to reintroduce BASL expression in a cell cycle-dependent manner during stomatal lineage ACDs (Figure 2A). To restrict BASL expression to cells before division, we fused the BASL coding sequence to the destruction box (DBox) motif of CYCLINB1;1, which targets the fusion protein for degradation during anaphase (Genschik et al., 1998), and placed this cassette under the native BASL promoter. We will refer to this pre-division BASL reporter (*pBASL::Venus-DBox-BASL*) as “BASL^pre^”. To provide BASL primarily after division, we used the promoter of the cytokinesis-specific gene *KNOLLE* (Lauber et al., 1997) to drive fluorescently tagged BASL fused to a non-functional, mutated DBox (Genschik et al., 1998). This post-division BASL reporter (*pKNOLLE::Venus-mDBox-BASL*) will be referred to as “BASL^post^”. As a control, *BASL* was fused to the non-functional DBox and supplied in its complete expression range by *pBASL::Venus-mDBox-BASL*, referred to as “BASL^full^ “. Unless otherwise stated, all results reported below are from the expression of the indicated reporter in a *basl* -/- mutant background.

**Figure 2.**
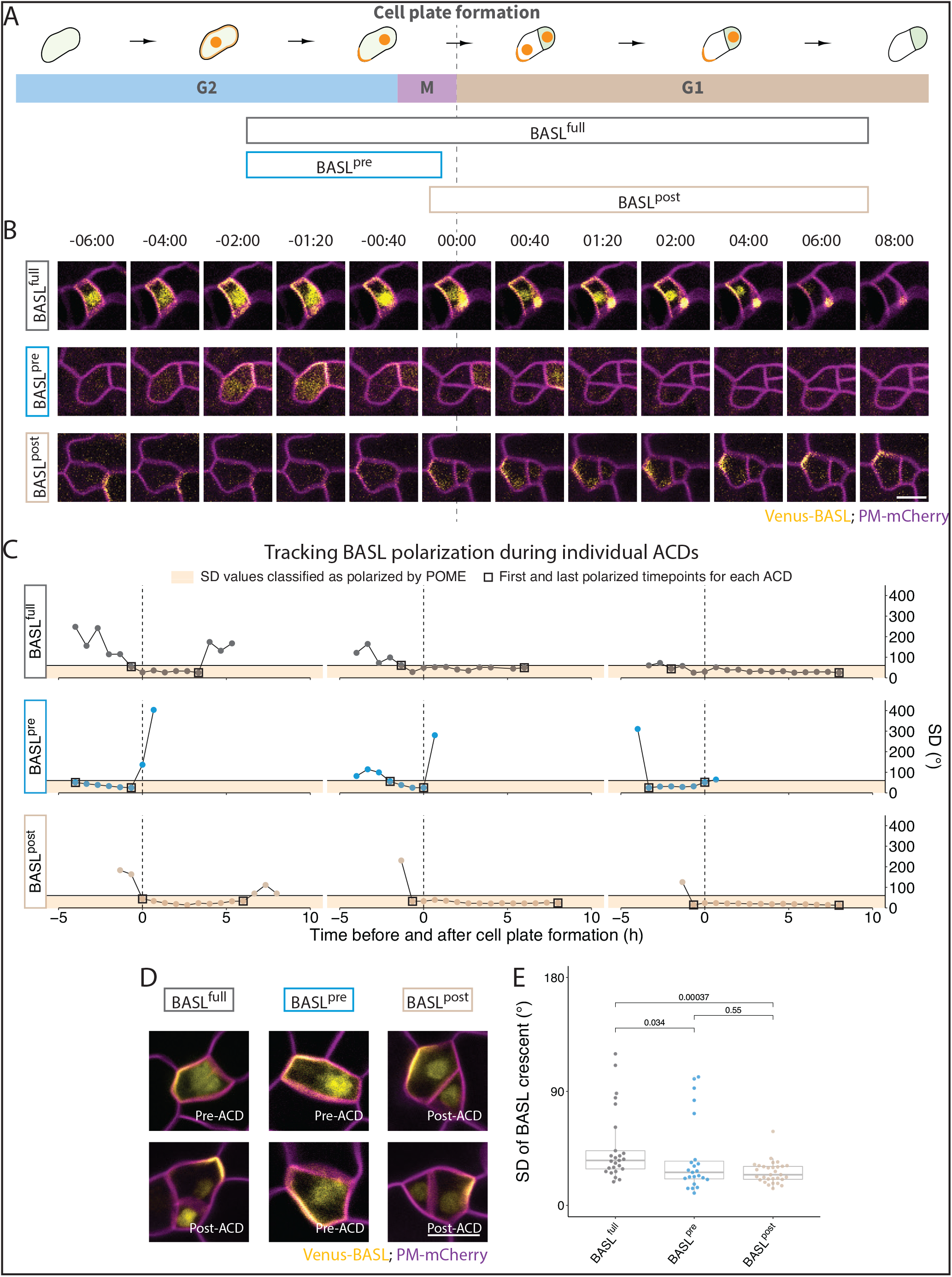
Genetic reagents to control BASL expression timing during stomatal lineage divisions. (A) Schematic illustrating the expression windows of BASL^full^, BASL^pre^, and BASL^post^ variants during asymmetric cell divisions. (B) Time-lapse images showing localization dynamics of BASL variants (yellow) during a single division in 3 dpg cotyledons. 00:00 (hours: minutes) marks cell plate formation. (C) Quantitative polarity tracking of BASL variants. Polarity crescent sizes are expressed as SD (°) in three individual cells of each genotype and were tracked using POME ((Gong, Varnau, et al., 2021), see methods); each circle represents a timepoint and squares mark the first and last points below the SD value of 60°, used as the cut-off for polarization. Vertical dashed lines indicate the frames during which the cell plate forms, and all values in the shaded orange zone are classified as polarized. (D-E) Comparison of crescent sizes among BASL variants, with two examples of each BASL variant (yellow) shown in D, and POME quantification of BASL crescent size (SD) of >20 cells/ variant shown in (E). In (E), the Bonferroni corrected p-value threshold = 0.017. In (B) and (D) cell outlines (magenta) are visualized by the plasma membrane reporter *pATML1::RCI2A-mCherry*. All p-values are calculated by Mann-Whitney test. Scale bars, 10 μm.

To test whether our temporally-restricted BASL variants recapitulated typical polarity dynamics, we performed time-lapse imaging to monitor the expression patterns of BASL^full^, BASL^pre^, and BASL^post^ during stomatal lineage ACDs (Figure 2B-C). In these time-lapse studies, BASL^full^ polarity dynamics were similar to previously published functional translational reporters (Dong et al., 2009; Gong, Varnau, et al., 2021). BASL^full^ was polarized at the cell cortex a few hours before cytokinesis (visualized by cell plate formation) and stayed polarized for more than 6 hours after the division (Figure 2B-C, top rows). BASL^pre^ was polarized only prior to cytokinesis, and BASL^pre^ signal was rapidly degraded (within one frame) after completion of cytokinesis (Figure 2B-C, middle rows). In contrast, BASL^post^ was detected during cytokinesis (00:00) or one frame (40 mins) earlier (−00:40). BASL^post^ expression and polarity became more apparent several hours after division and persisted for a similar amount of time as BASL^full^ (Figure 2B-C, bottom rows). Tracking BASL variants in individual cells revealed some variation in the duration of polarization (Figure 2C), but overall, the BASL^pre^ exhibited the same expression levels and behaviors of BASL^full^ (and previously published BASL reporters) before division, as did BASL^post^ after division. Notably, BASL^post^ could be highly polarized after division even when the reporter showed no polarity before division (Figure 2C-D). BASL variants were also correctly oriented relative to landmarks like the nascent division plane, and each also accumulated in the nucleus (Figure 2D).

To add more precision to the designation of “polarized”, we used the polarity quantification tool POME to measure the proportion of cell circumference occupied by the BASL reporters (Gong, Varnau, et al., 2021). To account for the fact that BASL^pre^ is expressed in meristemoids and BASL^post^ is expressed in SLGCs, we normalized polarity measurements as much as possible by analyzing cells of similar sizes. All three BASL variants could be highly polarized (Figure 2E). Some less polarized cells were observed in BASL^pre^ and BASL^full^ reporter lines (outliers in 2E) and these likely correspond to meristemoids that have just initiated polarization, as has been documented before (Gong, Varnau, et al., 2021).

### BASL expression before division fails to rescue many aspects of asymmetric division

Having validated our temporally restricted BASL constructs, we tested whether BASL^pre^ and BASL^post^ could rescue the *basl* mutant phenotypes. First, we quantified the frequency of small cell clusters on the abaxial epidermis of 5 dpg cotyledons (Figure 3A-B) and the frequency of stomatal clusters on the abaxial epidermis of 14 dpg cotyledons (Figure 3C). BASL^full^ completely rescued both phenotypes and resulted in a normally patterned epidermis (Figure 3A), confirming that the addition of the mDBox motif did not interfere with BASL’s function. BASL^pre^ largely failed to rescue, with cotyledons exhibiting numerous small cell and stomatal clusters. BASL^post^, in contrast, mostly rescued stomatal clusters and partly rescued the small cell phenotype (Figure 3A-C). Importantly, simultaneously introducing both BASL^pre^ and BASL^post^ into *basl* fully rescued the mutant phenotypes (Figure 3A-C), supporting the idea that restricted temporal expression of BASL revealed discrete functions before and after division.

**Figure 3.**
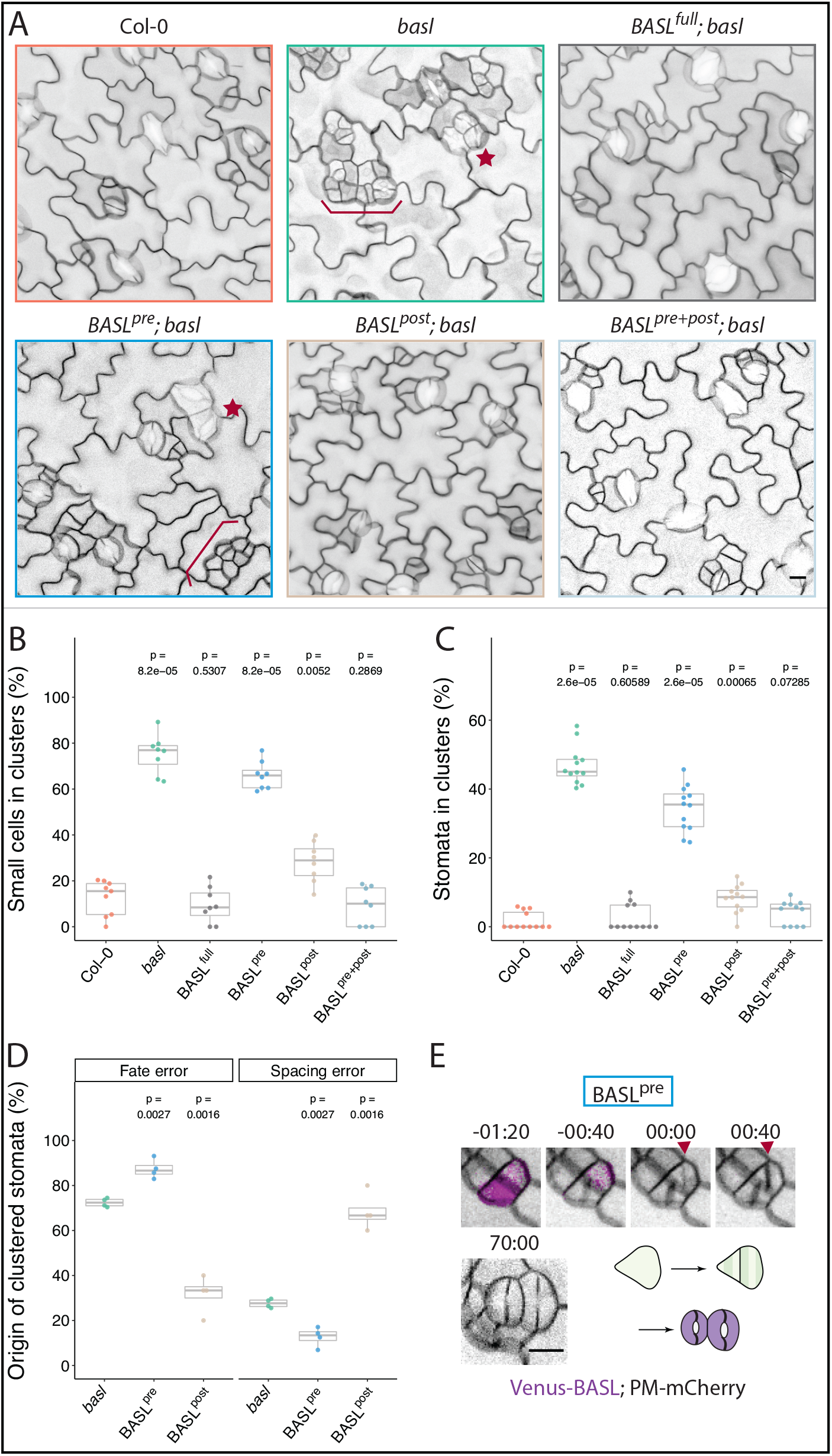
Temporally restricted BASL variants exhibit different capacities to rescue asymmetric defects in basl mutants. (A) Confocal images of 5 dpg abaxial cotyledon epidermis of Col-0, *basl*, and BASL variants rescuing *basl*. Braces and stars indicate clustered small cells and stomata, respectively. Cell outlines are visualized by plasma membrane reporters (magenta) *p35S::PIP2A-RFP* (*basl*) or *pATML1::RCI2A-mCherry* (all other lines). (B) Quantification of the clustered small cell phenotype in 5 dpg abaxial cotyledon epidermis of genotypes shown in (A) (n=8 388*388 μm2 fields/ genotype, Bonferroni corrected p-value threshold=0.01). (C) Quantification of the clustered stomata phenotypes in 14 dpg abaxial cotyledon epidermis of lines shown in (A) (n=12 623*467 μm2 fields/genotype, Bonferroni corrected p-value threshold=0.01).(D) Quantification of the origins of clustered stomata; all stomatal clusters formed from 3 dpg to 5 dpg were classified as originating from a fate or spacing error (n=4 cotyledons/ genotype; n>500 stomatal lineage cells).(E) Example of incorrect fate segregation in cells where BASL is only expressed before division (BASL^pre^, magenta). Cell outlines are visualized by the plasma membrane reporter (gray) *pATML1::RCI2A-mCherry*. Times denote (hours:minutes) with cell plate formation at 00:00. The p-values in (B-C) are calculated by Mann-Whitney test and the p-values in (B-C) are calculated by Student’s *t*-test due to small sample sizes. Pairwise p-values comparing individual genotypes to Col-0 (B-C) or *basl* (D) are presented.

These results suggested that not only was there a clear function for BASL after ACD, but that the post-divisional role might be more important than the pre-divisional role in defining stomatal lineage cell fates and patterning the leaf epidermis. Therefore, we performed a finer-grained phenotypic analysis to assay how BASL^pre^ and BASL^post^ affect cellular behavior. As shown in Figure 1G-H, stomatal clusters in *basl* had two origins: 1) a failure to establish differential fates in two sister cells and 2) misoriented spacing divisions. In BASL^pre^, clustered stomata were mostly derived from fate errors, where both sisters divided symmetrically to create paired guard cells (Figure 3D). In contrast, while the stomatal cluster phenotype was largely rescued by BASL^post^ (Figure 3C), the residual clustered stomata in BASL^post^ arose from spacing errors, where SLGC spacing divisions were misoriented, creating new stomata next to existing ones (Figure 3D and Supplemental Figure 1). Thus, we observed a clear divergence in the origin of clusters when comparing BASL^pre^ and BASL^post^. These results indicate that BASL contributes to different cellular and developmental processes before and after division, with pre-ACD BASL likely to regulate the orientation of ACDs, and post-ACD BASL enforcing cell fate differences in daughter cells. Indeed, extended time-lapse imaging and lineage tracing of lines lacking post-ACD BASL confirmed that cells undergoing morphologically normal ACDs can fail to correctly separate fates (Figure 3E).

### Presence of BASL before division coordinates crescent inheritance with daughter cell size asymmetry

In BASL^post^, the BASL crescent sometimes appears in the smaller daughter cell (Figure 4A, bottom right), a situation rarely seen in wildtype divisions. This gave us the unique opportunity to test whether BASL directly regulated cell size asymmetry, whether it coordinated size asymmetry with crescent inheritance, and ultimately whether cell size or crescent inheritance was the dominant factor in subsequent cell behavior. By quantifying the ratio of the surface area of the meristemoid to that of the SLGC in the frame following division (Figure 4B), we found that meristemoids were typically half the area of SLGCs in wildtype and in BASL^pre^. In BASL^post^ plants, however, meristemoid and SLGC sizes were more equalized, and we repeatedly identified cell pairs with “reversed” polarity, where the BASL crescent polarized in the smaller cell. Lineage tracing revealed that BASL presence was a better predictor of subsequent behavior than cell size, as the daughter cell with the BASL crescent, regardless of its size relative to its sister, always acquired the SLGC fate. Interestingly, the reversed cell size asymmetry also had consequences for cell lacking the BASL crescent. These large “meristemoids” underwent subsequent amplifying divisions more frequently than meristemoids produced from divisions with normal meristemoid/SLGC size asymmetry (Figure 4C). Thus, polarized BASL has a dominant impact on cell fate establishment, while cell size appears to influence the long term self-renewing capacity of stomatal lineage stem cells.

See also Supplemental Figure 1: Quantification of absolute numbers of stomatal clusters in BASL variant lines

**Figure 4.**
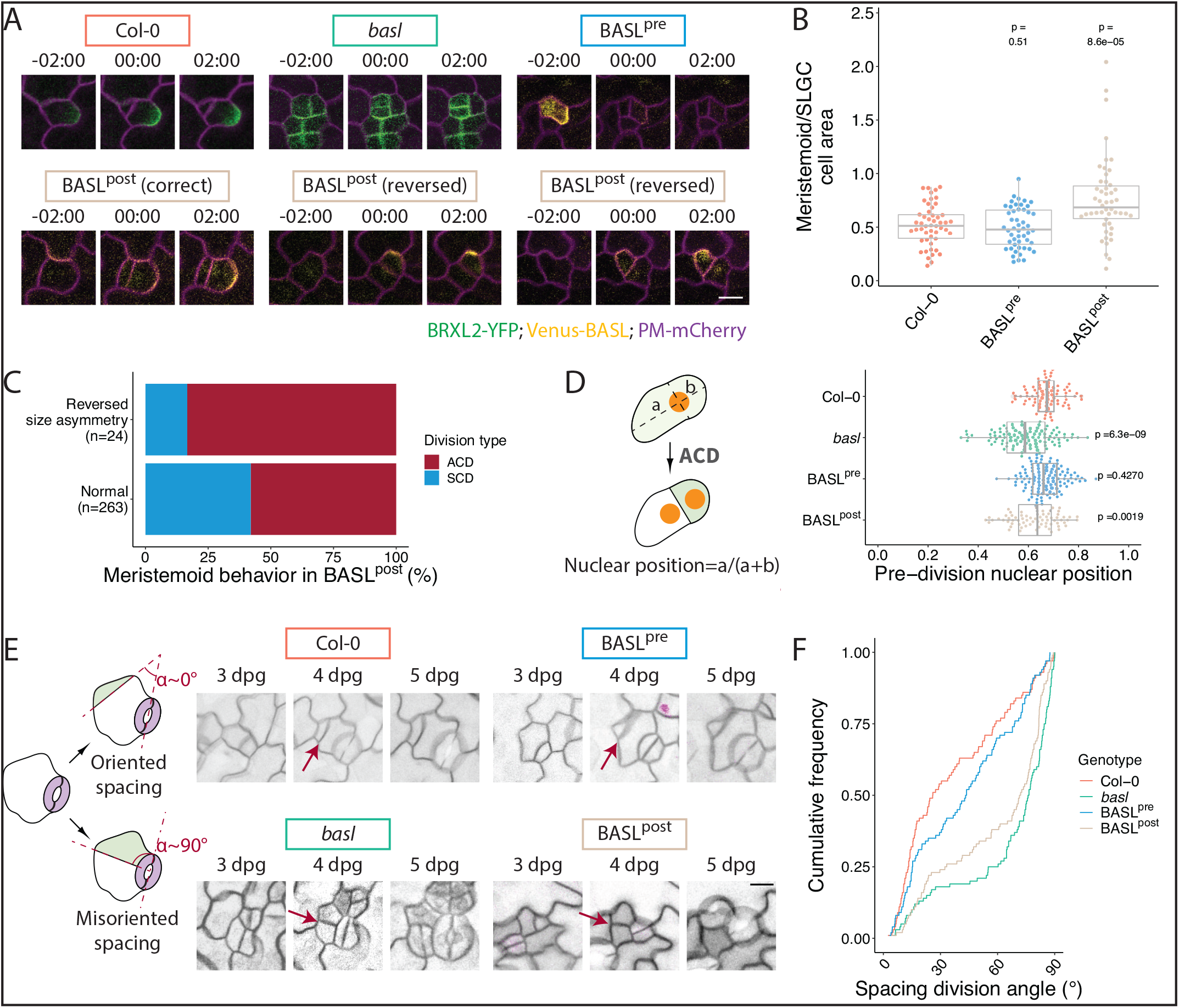
Presence of BASL before division influences division plane placement and coordinates crescent inheritance with cell size asymmetry. (A) Time-lapse images of ACDs in 3 dpg cotyledons of Col-0, *basl*, and BASL variants. Cell outlines are visualized by the plasma membrane reporter (magenta) *pATML1::RCI2A-mCherry* or *35S::PIP2A-RFP*. 00:00 (hours: minutes) marks cell plate formation. Polarity domains are labelled with *pBRXL2::BRXL2-YFP* (green, in Col-0 and *basl*) or by BASL variants (yellow). Examples of BASL^post^ include two divisions where the smaller daughter cell inherits the BASL crescent (reversed). (B) Quantification of cell size asymmetries from genotypes in (A), excluding *basl* because relationships between daughter cell size and crescent inheritance cannot be assigned (n=50 cells/genotype). (C) Quantification of meristemoid fates in BASL^post^ following divisions that exhibit normal and reversed size asymmetries. (D) Quantification of the final nuclear position pre-division in the indicated genotypes (n>70 cells/genotype, Bonferroni corrected p-value threshold=0.017). (E) Examples of spacing division orientations; α represents the angle between the stomatal pore axis and new meristemoid. Red arrows point to examples of scored divisions. (F) Graph of the cumulative frequencies of spacing division angles (from 0° to 90°) for each genotype in (E) (n=100 cells/genotype). All p-values are calculated by Mann-Whitney test and all pairwise p-values comparing individual genotypes to Col-0 are presented. Scale bars, 10 μm.

### BASL expression before division directs nuclear migration to create cell size asymmetries and to orient spacing divisions

Of the events comprising an ACD, we expected division plane placement to require pre-division polarity. Recent work has linked division plane position to BASL-directed nuclear migrations in the stomatal lineage (Muroyama et al., 2020). To test whether the reduced cell size asymmetry could be due to nuclear migration defects, we introduced a nuclear marker (*ML1::H2B-YFP*) into the BASL variant lines and characterized nuclear movements with time-lapse analysis. Pre-ACD nuclear migration was impaired in BASL^post^ and *basl*, as evidenced by the more centered position of the nucleus just prior to cell division (Figure 4D). This failure in nuclear migration provides an explanation for how cell size can be uncoupled from the presence of polar crescent in BASL^post^ plants.

Orienting division planes to preserve or avoid cell contacts may be an even more critical role for pre-ACD BASL than regulating cell size asymmetry in the stomatal lineage. During spacing divisions, the division plane is placed away from the existing stoma or stomatal precursor to avoid the formation of stomata pairs (Figure 4E, left), and this oriented division pattern is dependent on BASL activity (Figure 1H, 3D). We reasoned that BASL’s ability to orient spacing divisions relies mostly on its ability to direct pre-divisional nuclear migration. This hypothesis was supported by lineage tracing experiments where we observed more misoriented spacing divisions in the cotyledons of *basl* and BASL^post^ plants (Figure 4E). Quantification of spacing division angle in these genetic backgrounds revealed that without pre-ACD BASL activity, the SLGC’s ability to orient spacing division was compromised, and more spacing divisions were orthogonal to existing stomata (Figure 4F). Many cases of such misoriented spacing divisions led to the formation of stomatal pairs (Figure 4E, right).

### Presence of BASL after division coordinates division propensity between sister cells resulting from ACDs

A prominent phenotype in BASL^pre^, in addition to the stomatal clusters arising from cell fate defects, was an excessive number of clustered small cells (Figure 3B). The arrangement of cells in these clusters suggested that both daughters of an ACD divided more frequently than in wildtype (Figure 5A). This phenotype was also prominent in *basl* but not in BASL^post^ (Figure 5A). In lineage traces of cells in the wildtype Col-0 cotyledon epidermis (followed from in 3dpg to 5dpg), we noticed that spacing divisions were less frequent in SLGCs whose sister meristemoid underwent an amplifying division and more frequent in SLGCs whose sister meristemoid differentiated into a GMC (and ultimately became guard cells) (Figure 5B). This suggested some type of coordination between the division propensity of sister cells. We quantified the extent of potential coordination by tracking how often both a spacing division and an amplifying division occurred in a pair of sister cells (Obs[SPA+AMP]) and comparing this to the expected frequency if spacing and amplifying divisions were independent events (Exp[SPA+AMP], see details in methods). We found a large difference between expected and observed frequencies in wildtype Col-0 (Figure 5C). To test whether this presumed coordination between sisters was robust to increases in overall division frequency we quantified divisions in plants expressing an additional copy of the transcription factor SPEECHLESS (*SPCHpro::SPCH-CFP* or SPCH +) (Davies & Bergmann, 2014). Although both types of ACDs were elevated in SPCH + (Figure 5B), SLGC spacing division propensity was still highly correlated with the division behavior of its sister (Figure 5B-C). Interestingly, the coordination between sister cell divisions was diminished in *basl* and BASL^pre^ plants but maintained in BASL^post^ plants (Figure 5C). This suggests that meristemoid/SLGC sister cell pairs communicate to coordinate spacing and amplifying divisions via BASL activity post-ACD.

**Figure 5.**
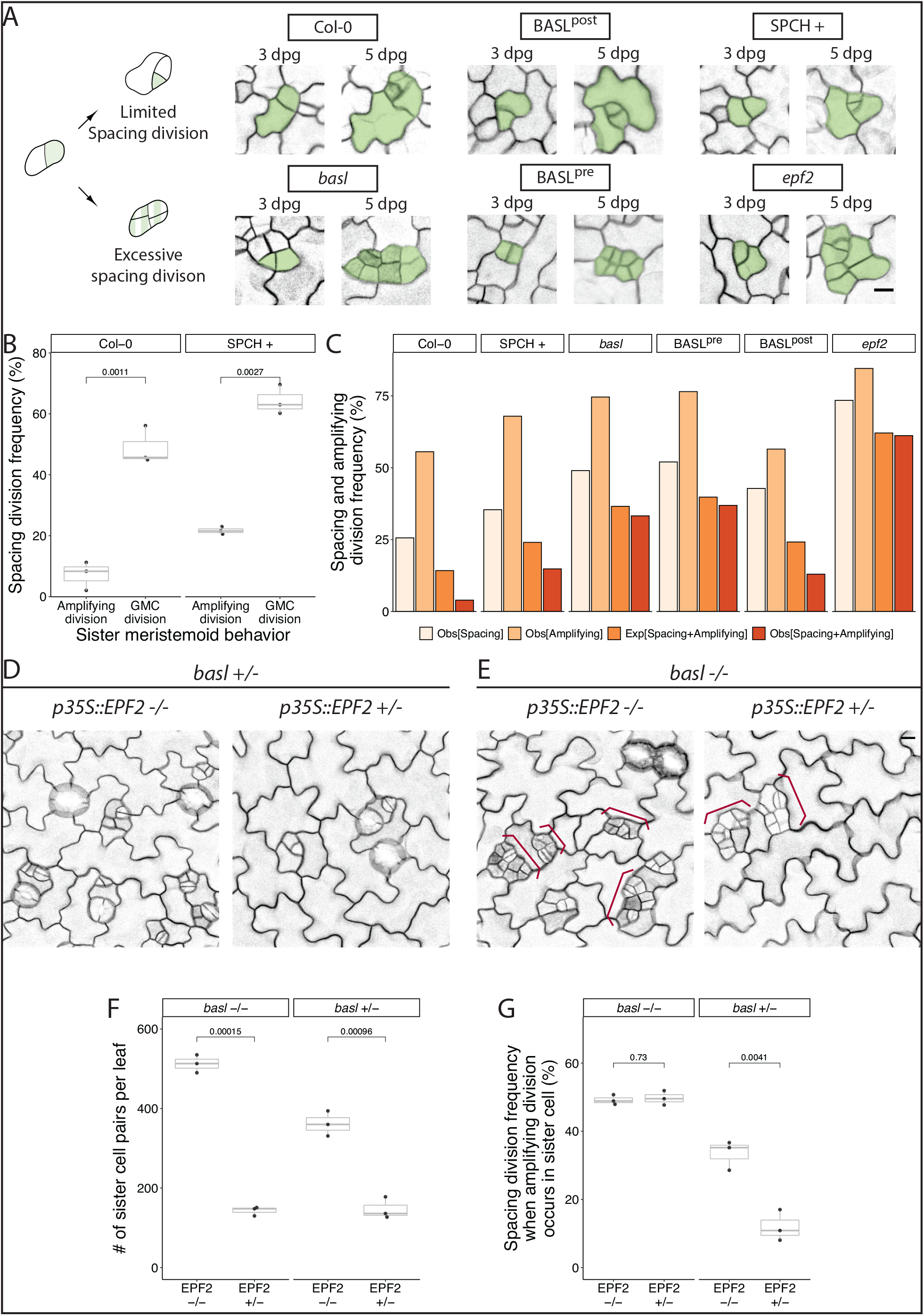
Presence of BASL post-ACD ensures cell fate asymmetry and coordinates sister cell divisions. (A) Schematic and examples of how excessive spacing divisions form small cell clusters. Top row gives examples of normal lineages where meristemoid amplifying divisions are associated with limited SLGC spacing divisions. Bottom row shows examples where both amplifying and spacing divisions occur in neighboring sisters. Tracked stomatal lineage cells are false colored in green. (B) Division frequencies are coordinated between sister cells when overall divisions are elevated through increased SPCH dosage (SPCH+). Frequencies are calculated by tracing the subsequent division behaviors of the daughters of an asymmetric division (n=3 cotyledons/genotype; n>300 pairs/cotyledons tracked from 3-5dpg) (C) Quantification of observed spacing division frequency (Obs[SPA]), observed amplifying division frequency (Obs[AMP]) and observed frequency of both amplifying division and spacing division (Obs[SPA+AMP]) compared with the expected frequency of amplifying division and spacing division (Exp[SPA+AMP]) if these two divisions were independent (calculated by multiplying Obs[AMP] by Obs[SPA]; n>500 pairs/genotype). (D) Suppression of entry and spacing divisions by overexpression of EPF2 (*35S::EPF2*) in BASL +/-plants. (E) Suppression of entry divisions, but not spacing divisions by *35S::EPF2* in *basl* -/-. (F-G) Quantification of entry divisions (F) estimated by the number of actively dividing pairs of sister cells at 3dpg) and observed spacing divisions (G) from genotypes shown in (D-E). Both frequencies are quantified from whole leaf lineage tracing experiments (n=3 cotyledons/genotype tracked from 3-5dpg; n>120 pairs/cotyledons in *35S::EPF2* lines; n>300 pairs/cotyledons in lines not expressing *35S::EPF2*). In (D-E), cell outlines are visualized by the plasma membrane reporter *pATML1::RCI2A-mCherry* and small cell clusters are labeled with red braces. All p-values are calculated by Student’s *t*-test due to small sample size. Scale bars, 10 μm.

How might BASL help coordinate cell division frequency between stomatal lineage sister cells? One hypothesis is that the peptide ligand EPIDERMAL PATTERNING FACTOR2 (EPF2), produced by meristemoids, signals to its sister SLGC. EPF2 was previously shown to suppress ACDs in neighbor cells through activating MITOGEN ACTIVATED PROTEIN KINASE (MAPK) signaling (Han & Torii, 2016; Hara et al., 2009; Hunt & Gray, 2009). MAPK signaling, in turn, is facilitated by BASL-mediated scaffolding post-ACD (Zhang et al., 2015). We re-examined the *epf2* loss of function mutant and found that, like *basl* and BASL^pre^, the phenotype included excessive spacing divisions and a lack of sister cell division coordination (Figure 5A, C). We then asked whether BASL was required for this EPF2-dependent inhibition of spacing divisions. Overexpression of EPF2 has a dose-dependent effect on ACDs (Lee et al., 2015) and by expressing a *p35S::EPF2* construct, we were able to generate lines where most, but not all, divisions were reduced. By crossing such a *p35S::EPF2* line made in a basl background to basl and Col-0 plants and assaying phenotypes in the F1 generation, we tested whether BASL mediated the reduction in divisions. Overexpression of EPF2 suppressed both stomatal entry divisions and spacing divisions in *BASL* +/-(Figure 5D, F, and G), while overexpression of EPF2 in *basl* -/-suppressed only entry divisions but not the excessive spacing divisions (Figure 5E-G). *BASL* is therefore required for EPF2-driven division suppression, and EPF2 appears to be the signal coordinating the trade-off between division propensity of meristemoids and their sister SLGCs.

## DISCUSSION

Polarity domains can act as hubs to coordinate multiple cellular behaviors across the cell cycle. Our understanding of how these potentially independent functions are coordinated in plant cells, however, has been limited by the resolution of traditional mutant phenotype analysis. Here, we developed new experimental approaches for the quantitative analysis of developmental dynamics of the *Arabidopsis* stomatal lineage, enabling new insights into the rules that govern cell behaviors and tissue patterning. Our temporal dissection of BASL function confirmed previous models implicating *BASL* in the regulation of nuclear migration before division and reinforcement of cell fate after division (Dong et al., 2009; Muroyama et al., 2020; Zhang et al., 2016). In addition, our data extend previous models of BASL function in two significant ways. First, our BASL^post^ data suggest that the critical activity of polarized BASL does not need to be inherited through the division but can function when formed de novo after ACD. Second, our system allowed us to uncouple cell size and polarity status, which are typically highly correlated in stomatal lineage divisions.

The post-division activities of BASL are particularly interesting because most of the current literature on polarity proteins during ACDs focuses on their roles in regulating cellular processes such as spindle orientation and fate determinant segregation before division. The *Arabidopsis* stomatal lineage, with its long cell cycles and incorporation of cell growth between divisions, contrasts with well-studied ACD models like *C. elegans* embryos or *Drosophila* neuroblasts but is representative of the typical situation in plant (and in mammalian) development (Francis et al., 2008; Gonsalvez et al., 2015; Jedrusik et al., 2010). The flexibility of the stomatal lineage, where the fate of cells that undergo ACDs are not predetermined, may require different mechanisms or layers of regulation than systems with invariant developmental trajectories. Building from the observation that, during the long cell divisions of the *Arabidopsis* stomatal lineage, polar cortical domains persist for hours after division (Gong, Alassimone, et al., 2021; Gong, Varnau, et al., 2021), we had an opportunity to use genetic tools to supply BASL before and after division to dissect its roles at those stages. We concluded that the ability of BASL to generate fate asymmetry in ACD daughter cells largely depends on its post-divisional polarity, a different result from classical ACD systems; for example, in *Drosophila* neuroblasts, the differential inheritance of fate determinant factors such as Numb, Prospero and Brat, depends on PAR polarity prior to ACD (Homem & Knoblich, 2012). We have several hypotheses for why the stomatal lineage appears to emphasize post-divisional activities: (1) in a flexible system, it may be that segregated determinants bias a cell fate decision but can be overridden by signaling or other events post division or (2) polarized BASL and the signaling molecules it scaffolds are themselves the fate determining factors. In either case, post-ACD BASL is required to maintain the separate cell fates.

Another interesting comparison between animal models and the stomatal lineage is the effect of cell size on asymmetric divisions. Divisions creating daughters of unequal sizes are common during development, but whether cell size *per se* matters for fate is not clear. Recent work experimentally equalizing cell size in *C. elegans* embryos showed effects on cell cycle timing and on developmental progression (Jankele et al., 2021). In our work, we found that cell size did not correlate with cell divisions in the same way; rather than larger cells dividing faster as in *C. elegans*, small cells of the stomatal lineage divided faster, though loss of polarity proteins equalized timings in both situations. In terms of fate, smaller daughters of an ACD that expressed BASL^post^ (in the case of ACD size asymmetry reversal) became pavement cells, suggesting that size was not critical for cell identity during stomatal lineage ACDs. Cell size, however, did affect the number of divisions the non-BASL inheriting cell underwent, suggesting that in this flexible stem-cell like lineage, cell size asymmetry may be important for regulation of overall division capacity.

Investigation of the small cell cluster phenotype was particularly fruitful for refining and defining new roles for BASL. First, we found that *BASL* was haploinsufficient for this phenotype, suggesting that the mechanisms regulating cell division propensity are more sensitive to BASL dosage than those that are required for cell fate. Because the stomatal clustering phenotype is completely recessive, however, it indicates that the small cell clusters can be resolved, and this suggests additional layers of regulation that enable error correction. Second, lineage tracing of cells in small cell clusters revealed a previously unappreciated coordination between the division behaviors of meristemoids and their sister SLGCs. Our data indicate that BASL is an effector of EPF2-mediated signaling between sister cells that prevents both cells from undergoing ACDs. This finding forces us to consider the idea that polarity domains can have non-cell autonomous effects; future studies might determine whether this mechanism occurs in other contexts and whether the non-autonomous effects are chemical (signaling) or mechanical in nature. Why might stomatal lineage sister cells coordinate their divisions? Coordination could serve as patterning mechanism to prevent divisions that would place guard cell precursors in contact. Alternatively, plants alter stomatal indices (SI, the ratio of stomatal to non-stomatal epidermal cells) in response to systemic and environmental cues. However, amplifying and spacing divisions have opposite effects on SI, with amplifying divisions decreasing, and spacing divisions increasing SI. The antagonistic behaviors of sisters may reflect circuitry designed to amplify cues that instruct the tissue to modulate SI in response to environmental stimuli.

## MATERIALS AND METHODS

### Plant material and growth conditions

All *Arabidopsis* lines used in this study are in Col-0 background. All *Arabidopsis* seeds were surface sterilized by bleach or 75% ethanol and stratified for 2 days. After stratification, seedlings were vertically grown on 1/2 Murashige and Skoog (MS) media with 1% agar for 3-14 days under long-day condition (16h light/8h dark at 22 °C) and moderate intensity full-spectrum light (110 μE).

Previously reported mutants and transgenic lines include: *epf2* (Hunt & Gray, 2009), *pATML1::RCI2A-mCherry* (Bringmann & Bergmann, 2017), *p35S::PIP2A-RFP* in *basl-2* (Rowe et al., 2019), *pATML1::H2B-YFP p35S::PIP2A-RFP* in *basl-2* (Muroyama et al., 2020), and *pATML1::mCherry-RCI2A pRPS5A::DII-n3xVenus pRPS5A::mDII-ntdTomato pBRXL2::BRXL2-YFP* (Muroyama et al., 2020).

Newly generated lines include: *pBASL::Venus-mDBox-BASL pATML1::mCherry-RCI2A* in *basl-2, pBASL::Venus-DBox-BASL pATML1::mCherry-RCI2A* in *basl-2, pKNOLLE::Venus-mDBox-BASL pATML1::mCherry-RCI2A* in *basl-2*, and *p35S::EPF2 p35S::PIP2A-RFP* in *basl-2*. For nuclear migration assays, the BASL variants were crossed with *pATML1::H2B-YFP p35S::PIP2A-RFP* in *basl-2*, and plants were scored in F1s as the large number of transgenes lead to silencing in later generations. To ensure the same dose of EPF2 overexpression, *p35S::EPF2/+ p35S::PIP2A-RFP* in *basl-2* was crossed to either *pATML1::RCI2A-mCherry* (Col-0) or *p35S::PIP2A-RFP* in *basl-2* and phenotypic assays were performed in F1 seedlings.

### Vector construction and plant transformation

To generate pBASL::Venus-DBox-BASL, the coding sequence of the destruction box (DBox) domain of *Arabidopsis* CYCB1;1 (first 116 amino acids) (Genschik et al., 1998) was fused with the coding sequence of fluorescent protein Venus utilizing a BamHI restriction site, cloned into Gateway vector pENTR (Thermo Fisher), and recombined with entry vectors containing the *BASL* promoter (3kb) and the BASL coding sequence into binary vector pK7m34GW (Karimi et al., 2007). Similar approaches were used to generate *pBASL::Venus-mDBox-BASL*, and *pKNOLLE::Venus-mDBox-BASL* with the *BASL* and 2.4 kb *KNOLLE* promoter respectively. mDBox contain the same region of *Arabidopsis* CYCB1;1 as DBox, but with the motif RxxLxx(L/I)xN changed to GxxVxx(L/I)xN (Genschik et al., 1998). To generate *p35S::EPF2*, the *EPF2* coding sequence was cloned into pENTR and recombined into pH35GS binary vector (Kubo et al., 2005). Transgenic plants were then generated by Agrobacterium-mediated transformation (Clough & Bent, 1998), and transgenic seedlings were selected on 1/2 MS plates with 50 μg/mL hygromycin (pH35GS) or 50 μg/mL kanamycin (pK7m34GW) antibiotic.

### Microscopy, image acquisition, and image analysis

All fluorescence imaging experiments were performed on a Leica SP5 confocal microscope with HyD detectors using 40x NA1.1 water objective with image size 1024*1024 and digital zoom from 1x to 2x.

For time-lapse experiments, 3 dpg seedlings were mounted in a custom imaging chamber (Davies & Bergmann, 2014) filled with 1/2 MS solution. For the time-lapse experiments reported in this study, there was a 30-45 min interval between each image stack capture. For the time-lapse followed by time course experiments, seedlings were imaged in the time-lapse chamber for 16 hours as above. After imaging, seedlings were removed from the imaging chamber and returned to MS-agar plates (with appropriate supplements) for 48 hours under standard light and temperatures. The same epidermal surface from the same plant was re-imaged again to capture the developmental outcome of all the epidermal cells. Whole leaf based time-course and lineage tracing experiments were performed as described in Gong, Alassimone, et al. (2021). Quantification of nuclear migration was done as in Muroyama et al. (2020).

Quantification of BASL polar crescent size was done with the FIJI plugin POME (Gong, Varnau, et al., 2021). 20-30 highly expressing cells were selected, and the BASL localization pattern in each of these cells was analyzed by POME and fitted to a Gaussian model to extract key parameters of BASL localization pattern. These parameters included standard deviation σ, center μ, amplitude α, and baseline value β. The standard deviation (σ) from the regression model of each cell’s BASL localization pattern was then defined as the crescent size. The crescent sizes of all analyzed cells from BASL variants were then plotted and compared.

To quantify the small cell cluster phenotype, seedlings bearing a fluorescent cell outline marker were imaged at 5 dpg with a 388*388 μm2 field of view. Each individual image was segmented in FIJI. Cells with surface area smaller than 120 μm2 were considered small cells. To simplify the analysis, we used cell area instead of cell volume since epidermal cell height (distance from apical to internal face) is quite uniform at the cotyledon ages we measured (Muroyama et al., 2020). Four or more small cells in contact were considered a cluster and the fraction of small cells in clusters/ total small cells counted. To count clusters of mature stomata, seedlings were collected at 14 dpg. Samples were cleared with 7:1 solution (7:1 ethanol:acetic acid), treated with 1N potassium hydroxide solution, rinsed in water, and then mounted in Hoyer’s solution.

Individual leaves were then imaged with a Leica DM6B microscope with 20x NA0.7 air objective in differential contrast interference mode. Total stomata number and the number of stomata in direct contact were counted to calculate the percentage of stomata in clusters. The origin of a given stomatal cluster was determined from lineage tracing of whole leaf time courses (3 dpg to 5 dpg). Stomata clusters generated from a pair of sister cell from a cell division was classified as fate error while any stomata cluster generated from a misplaced spacing division was classified as spacing error.

To determine whether sister cell division propensities were correlated, we tracked sister cell pairs for 48 hours (3-5 dpg). We then calculated the frequencies of spacing and/or amplifying divisions among tracked cells. The observed spacing division frequency Obs[SPA] is the number of SLGCs in which a spacing division occured divided by the total number of tracked SLGCs. The observed amplifying division frequency Obs[AMP] is the number of meristemoids in which an amplifying division occurred divided by the total number of tracked meristemoids. Obs[SPA+AMP] is the observed frequency of meristemoids undergoing an amplifying division and their sister SLGC undergoing a spacing division divided by the total number of tracked sister pairs. From the lineage tracing data, we also computed Exp[SPA+AMP], the expected frequency of a pair of sister cells undergoing an amplifying division and a spacing division if these two divisions were independent of each other, by multiplying Obs[SPA] and Obs[AMP]. Deviations of Obs[SPA+AMP] from Exp[SPA+AMP] suggest interactions between sisters, with a lower Obs[SPA+AMP] relative to Exp[SPA+AMP] indicative of competition or inhibition between sisters. The division behaviors of at least 500 cell pairs were tracked for each genotype.

### Statistical analysis

All statistical analyses were performed in RStudio. Unpaired Mann-Whitney tests and Student’s t-test were conducted to compare two data samples with compare_means function from the ggpubr package (Kassambara, 2020). For all graphs, p-values from the unpaired Mann-Whitney tests or Student’s t-test were directly labelled on these graphs. Bonferroni corrections were performed when more than 2 pair-wise comparisons were conducted to reduce Type I error. Bonferroni corrected p-value thresholds are indicated in the corresponding figure legends.

## Acknowledgments

We thank the members of the Bergmann lab for discussions and feedback on manuscripts and Chin-Min Kimmy Ho (now at IPMB, Academia Sinica, Taiwan) for constructs.

## Author contributions

Conceptualization: Y.G., J.A., D.C.B.; Methodology: Y.G., J.A., A.M.; Validation: Y.G., J.A., A.M., G.A., R.V., A. L.; Formal analysis: Y.G., A.M.; Investigation: Y.G., J.A., A.M., G.A., R.V., A. L.; Writing -original draft: Y.G., J.A., A.M., G.A., D.C.B.; Visualization: Y.G., A.M.; Supervision: D.C.B.; Project administration: D.C.B.; Funding acquisition: J.A., A.M., D.C.B.

## SUPPLEMENTARY INFORMATION

**Supplemental Figure 1.**
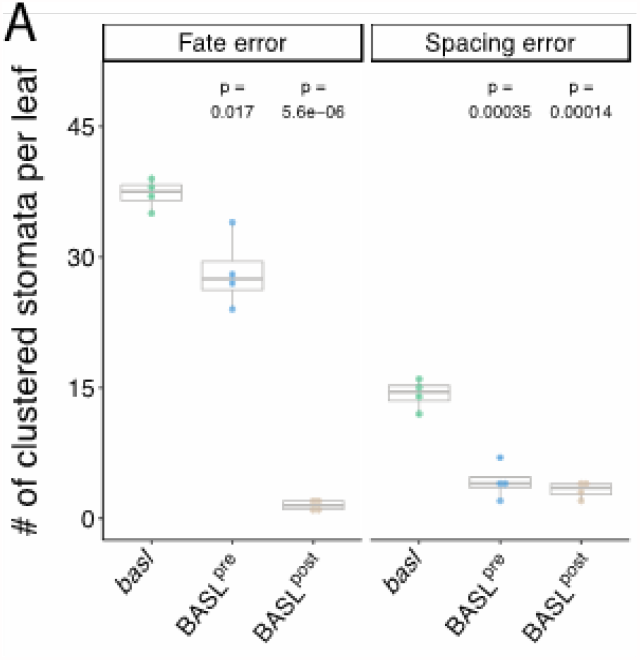
Quantification of absolute numbers of stomatal clusters in BASL variant lines. (A) Counts of stomatal clusters from which Figure 3D was calculated are calculated by Student’s *t*-test due to small sample sizes. Pairwise p-values comparing individual genotypes to *basl* are presented.

**Supplementary Table 1.**
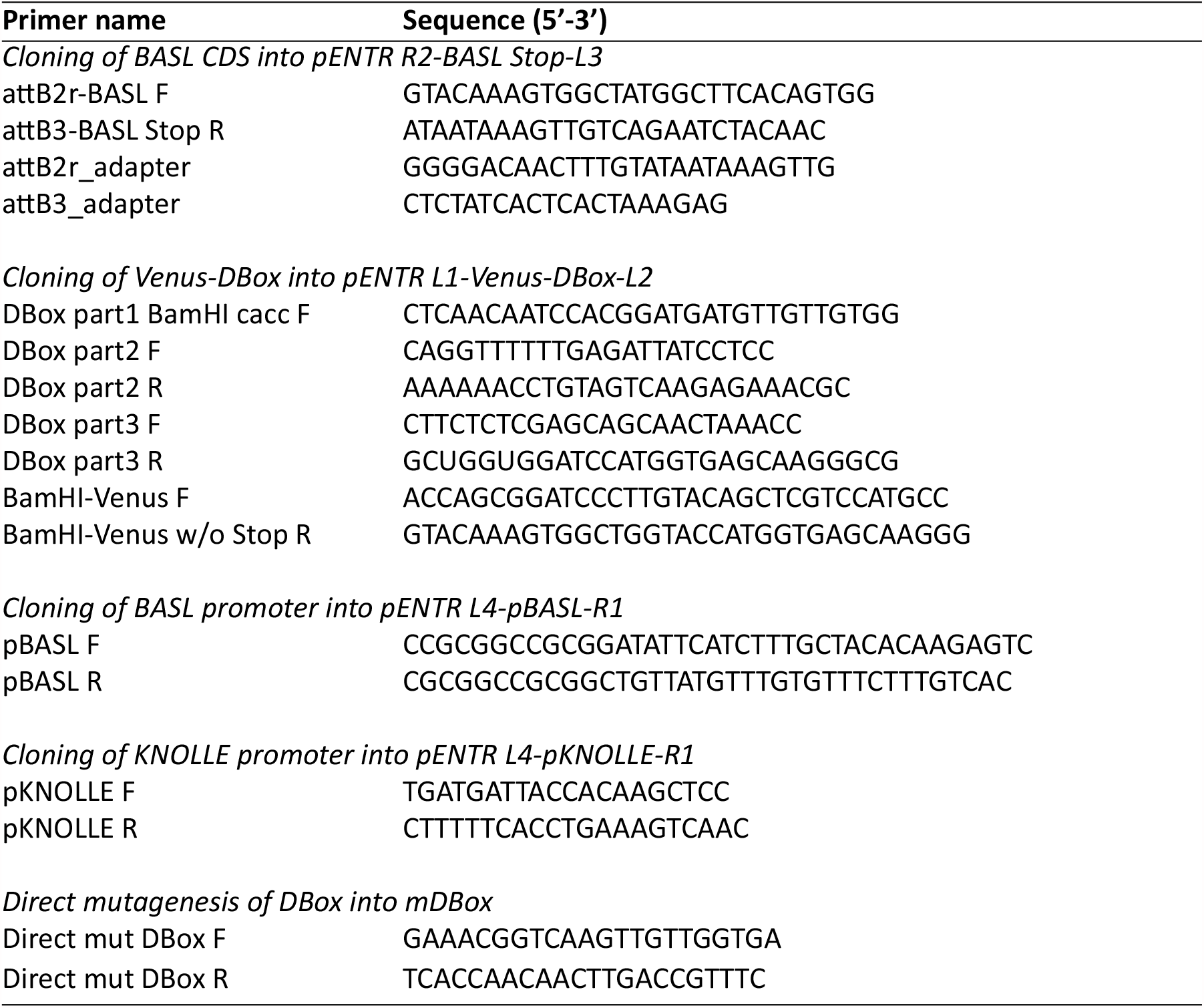
Primers used in this study.

